# Long-term temporal stability of peripheral blood DNA methylation alterations in patients with inflammatory bowel disease

**DOI:** 10.1101/2022.08.22.504377

**Authors:** Vincent Joustra, Andrew Y.F. Li Yim, Ishtu Hageman, Evgeni Levin, Alex Adams, Jack Satsangi, Wouter J. de Jonge, Peter Henneman, Geert D’Haens

## Abstract

**Introduction:** There is great current interest in the potential application of DNA methylation alterations in peripheral blood leukocytes (PBL) as biomarkers of susceptibility, progression and treatment response in inflammatory bowel disease (IBD). However, the intra-individual stability of PBL methylation in IBD has not been characterised. Here, we studied the long-term stability of all probes located on the Illumina HumanMethylation EPIC BeadChip array.

**Methods:** We followed a cohort of 46 adult IBD patients (36 Crohn’s disease (CD), 10 ulcerative colitis (UC), median age 44 (IQR: 27-56), 50% female) that received standard care without any intervention at the Amsterdam UMC. Paired PBL samples were collected at two time points with a median 7 (range: 2-9) years in between. Differential methylation and intra-class correlation (ICC) analyses were used to identify time-associated differences and temporally stable CpGs, respectively.

**Results:** Around 60% of all EPIC array loci presented poor intra-individual stability (ICC <0.50); 78.114 (≈9%) showed good (ICC 0.75 – 0.89); and 41.274 (≈5%) excellent (ICC ≥0.90) stability. Focusing on previously identified consistently differentially methylated positions indicated that 22 CD-, 11 UC-, and 24 IBD-associated loci demonstrated high stability (ICC ≥0.75) over time; of these, we observed a marked stability of CpG loci associated to the *HLA* genes.

**Conclusion:** Our data provide insight into the long-term stability of the PBL DNA methylome within an IBD context, facilitating the selection of biologically relevant and robust IBD-associated epigenetic biomarkers with increased potential for independent validation. These data also have potential implications in understanding disease pathogenesis.

## Introduction

Crohn’s disease (CD) and ulcerative colitis (UC) are chronic relapsing and remitting inflammatory bowel diseases (IBD) characterized by a wide variety of phenotypic manifestations^1^. While the aetiology of IBD remains unknown, it is thought to arise as a result of a complex interplay between the host and microbial composition, triggered by environmental factors, such as tobacco smoking or diet^2–4^.

Accordingly, much effort has been invested in understanding the interaction between host and environment, which is thought to be mediated by the epigenome^5^. The epigenome represents the set of mitotically heritable modifications that can affect gene transcriptions without altering the primary DNA sequence^6^. DNA methylation, one of the most studied epigenetic mechanisms, involves the attachment of methyl groups to cytosine-phosphate-guanine (CpG) nucleotide sequences on the DNA. This covalent attachment is mitotically heritable and can, under certain conditions, regulate gene expression thereby altering cellular behavior^7^. Over the past decade, multiple epigenome-wide association studies (EWAS) have sought to characterise, classify, and predict IBD and its various phenotypes using DNA methylation^8–13^. However, most EWAS in IBD to date have been cross-sectional in design, reporting aberrant DNA methylation signatures in peripheral blood leukocytes (PBL) and/or mucosal tissue^9,14–22^, with only a single longitudinal study in mucosal tissue^20^ and PBL^23^.

Previous literature has shown that the intra-individual variability of DNA methylation is most prominent during the early stages of life, which gradually diminishes and presents a more stable phenotype after 5 years of life^24, 25^. Nonetheless, the influence of ageing on genome-wide DNA methylation has been well-described in monozygotic twins^26, 27^ and unrelated healthy populations^28, 29^, demonstrating a global decrease in methylation as individuals age, as well as site-specific increases in methylation in CpG-rich areas, both of which are thought to result from dynamic external and internal environmental changes^30–32^. As epigenetics and thus DNA methylation, is cell-type specific, observed differences found in heterogeneous populations such as PBL or tissue might reflect differences in the cellular composition^33–35^. Nonetheless, age-related differences were found in more homogeneous populations, such as purified T-cells and monocytes^15, 36, 37^

Despite the strong effects of age on DNA methylation, a high correlation between baseline and follow-up methylation data in IBD pediatric mucosal tissue has been observed^20^. In contrast, IBD-associated differences in blood have shown to largely revert back to patterns observed in non-IBD controls during follow-up as the result of treatment and normalisation of CRP^23^. It is noteworthy that these studies focused only on a subset of IBD-associated CpGs, and did not report on the long-term stability of all CpG probes located on the Illumina HumanMethylation EPIC BeadChip array. While temporal stability and intra-individual variability in PBL-derived DNA methylation has been investigated in adult healthy individuals^38–41^ and patients with systemic lupus erythematosus (SLE)^42^, no such study has been conducted in IBD patients.

There is widespread interest in the application of epigenetic markers in personalization of treatment^43^. If epigenetic biomarkers are to be used as pathognomonic for IBD or its (sub)phenotypes, the features of interest would need to remain stable throughout the duration of the disease and hence over time without being affected by various internal/external exposures. In addition, for biomarker development, loci that are time-stable reduce the number of false positive findings thereby increasing the probability of independent replication. Furthermore, selection of time-stable epigenetic biomarkers would help overcome current practical barriers in sample collection at specific time frames thereby facilitating the use of larger samples sizes with similar phenotypes needed to enhance predictive power. We therefore sought to identify CpG positions that present stable DNA methylation in PBL obtained from a well-generalizable cohort of adult IBD patients with a 7-year median time between collected DNA samples.

## Methods

### Patient selection

We performed a single-centre, longitudinal EWAS, where we collected PBL samples from adult IBD patients at the Amsterdam University Medical Centers. The interval between the time of sampling ranged from 2 to 9 years with a median of 7 (Figure 1a). All included patients were historically diagnosed with either CD or UC on the basis of a combination of clinical symptoms and endoscopic inflammation as confirmed by histology per the current guidelines^44, 45^. In addition, all patients received standard care follow-up. No additional in- or exclusion criteria were used as the goal was to collect a cohort of IBD patients that reflected the overall IBD population at the Amsterdam University Medical Centers. This study was approved by the medical ethics committee of the academic medical hospital (METC NL24572.018.08 and NL53989.018.15) and written informed consent was obtained from all subjects prior to sampling.

**Figure 1.**
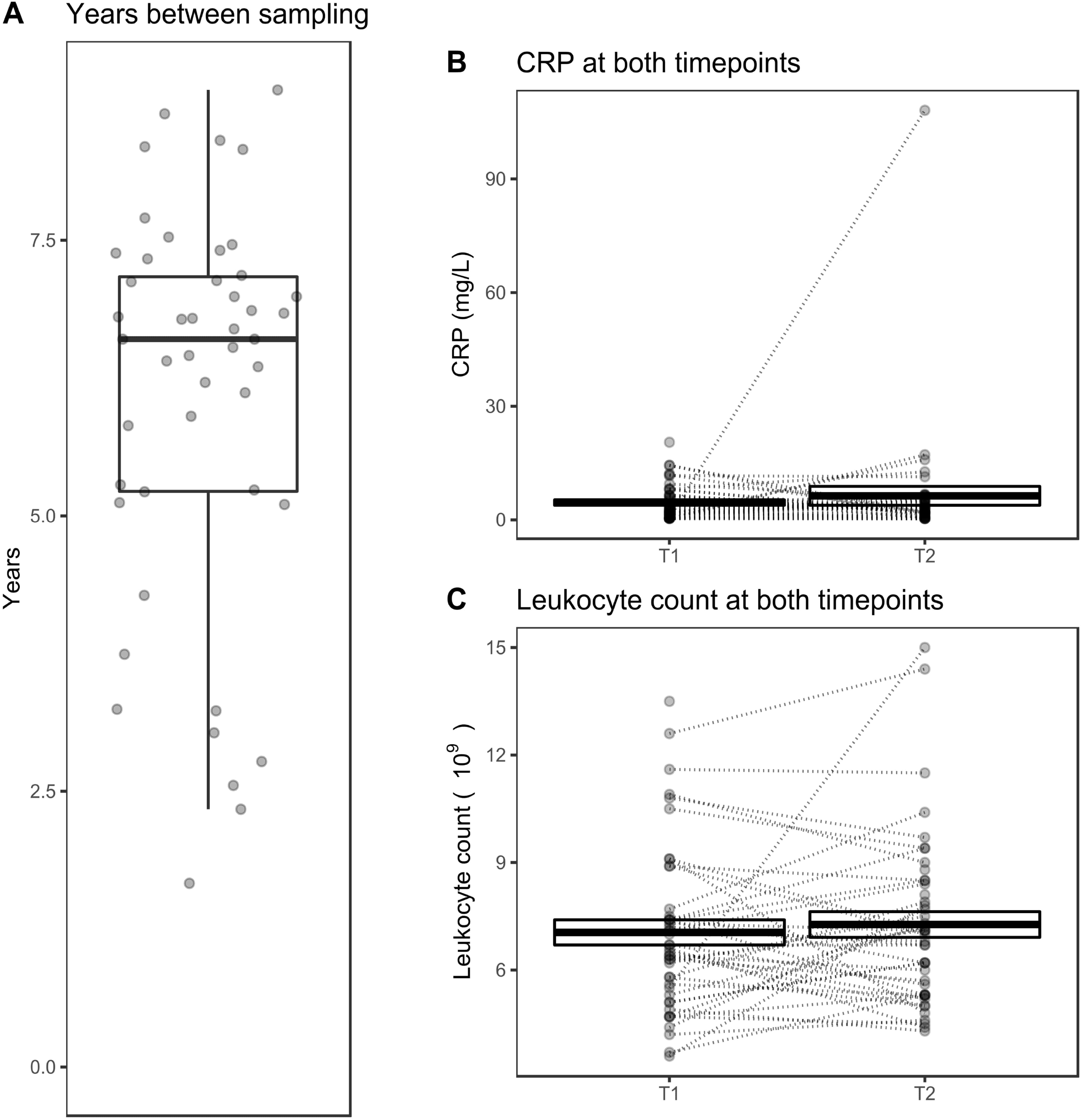
Patient characteristics over time. A) Visualisation of the number of years between both samplings per patient. Visualisation of the B) CRP (mg/L) and C) leukocyte count (10^9^) between both time points, where connected samples were obtained from the same patient annotated with the mean difference and *p*-value.

### Sample collection and DNA methylation analysis

Whole peripheral blood samples were collected in a 6mL EDTA tube and stored at −80 °C until further processing. Genomic DNA was isolated using the QIAsymphony whereupon the quantity of the DNA was assessed using the FLUOstar OMEGA and quality of the high-molecular weight DNA on a 0.8% agarose gel. Genomic DNA was bisulfite converted using the Zymo EZ DNA Methylation kit, randomized per plate to limit batch effects, and analysed on the Illumina HumanMethylation EPIC BeadChip array at the Core Facility Genomics, Amsterdam UMC, Amsterdam, the Netherlands.

### Statistical analysis of clinical data

Baseline characteristics of all included patients were summarised using descriptive statistics. Categorical variables are presented as percentages and continuous variables as median annotated with the interquartile range (IQR). Differences in CRP and leukocyte count levels between T1 and T2 were calculated using the Wilcoxon signed ranks test. Analyses of clinical data were performed in IBM SPSS statistics version 26 and methylation analyses in the R statistical environment version 4.2.1.

### Time dependent DNA methylation data analyses

For differential methylation analyses, raw DNA methylation data were imported into the R statistical environment using the Bioconductor *minfi^46^* package (version 1.36), whereupon the raw signal intensities were normalized using functional normalization^47^ and converted into methylation ratios. Differential methylation analyses was performed using *limma^48^* (version 3.46) and *eBayes^49^* regressing against time point (T2 vs T1), gender, smoking behaviour, disease, and blood cell distribution. Statistical significance was defined as a false discovery rate-adjusted *p*-value < 0.05. In addition to identifying time-associated differences in methylation, we also investigated differences in methylation associated with CRP and leukocyte count. Blood cell estimations were performed using the IDOL predictor CpGs as reference^50^. Time-associated DMPs were investigated for their association with age by performing gene set enrichment analyses using the age-associated CpGs reported by Horvath^51^, Hannum^52^, Levine^53^, and Knight^54^. Visualisations were generated using *ggplot2^55^* (version 3.3.5) and *gghighlight* (version 0.3.2).

### Time independent DNA methylation data analyses

For the time stability analyses, raw DNA methylation data were imported using *ewastools* to retain the 89 quality control probes that bind genetic variants (GVs). Methylation probes that might bind GVs were identified on the basis of the *minfi*-provided annotation files, which we termed as potential GVs.

Additional GV-binding probes were estimated using the Gaphunter tool^56^ as implemented in *minfi*, which we termed the predicted GVs. Moreover, as opposed to the differential methylation analysis, for the time stability analyses, we did not perform normalisation nor corrected for any other potential confounders (e.g. gender, smoking behaviour, disease, and blood cell distribution) in order to identify truly stable signals. Stability analyses were conducted using intraclass correlation (ICC) analyses, where ICC estimates and their 95% confidence intervals were calculated using the *irr* package implemented in R. Specifically, a two-way mixed, single measures, consistency analysis was performed^57^.

Visualisations were generated using *ggplot2^55^* (version 3.3.5) and *gghighlight* (version 0.3.2).

### GO enrichment and KEGG pathway enrichment analyses

Functional enrichment analyses genes annotated to both stable and unstable methylated probes was performed using GOmeth^58^ as implemented in *missMethyl^35^*. The GO terms were grouped according to biological process (BP), cellular component (CC) and molecular function (MF) and an FDR corrected p-value below 0.05 indicates a statistical significant difference.

### Data availability

The raw has been made available under controlled access in the European Genome-phenome Archive (accession ID: EGAS00001006501).

## Results

### Patient demographics

A total of 46 adult IBD (36 CD, 10 UC) patients with a median age of 44 (IQR: 27-56) years and median disease duration of 12 (IQR: 7-21) years were included. Gender, surgical history, disease location- and behaviour were balanced within this cohort (Table 1). Notably, 32 patients (69.6%) were previously treated with an anti-TNF, prior to T1 sampling. Between T1 and T2 during regular IBD care follow-up, 10 (21.7%) patients underwent IBD-related surgery, 24 (52.2%) were treated with vedolizumab and 14 (30.4%) with ustekinumab, reflecting the tertiary referral population seen at the Amsterdam UMC. No significant differences in median CRP (*p*-value = 0.97) or leukocyte count (*p*-value = 0.85) between T1 and T2 were observed (Figure 1b and 1c).

**Table 1.**
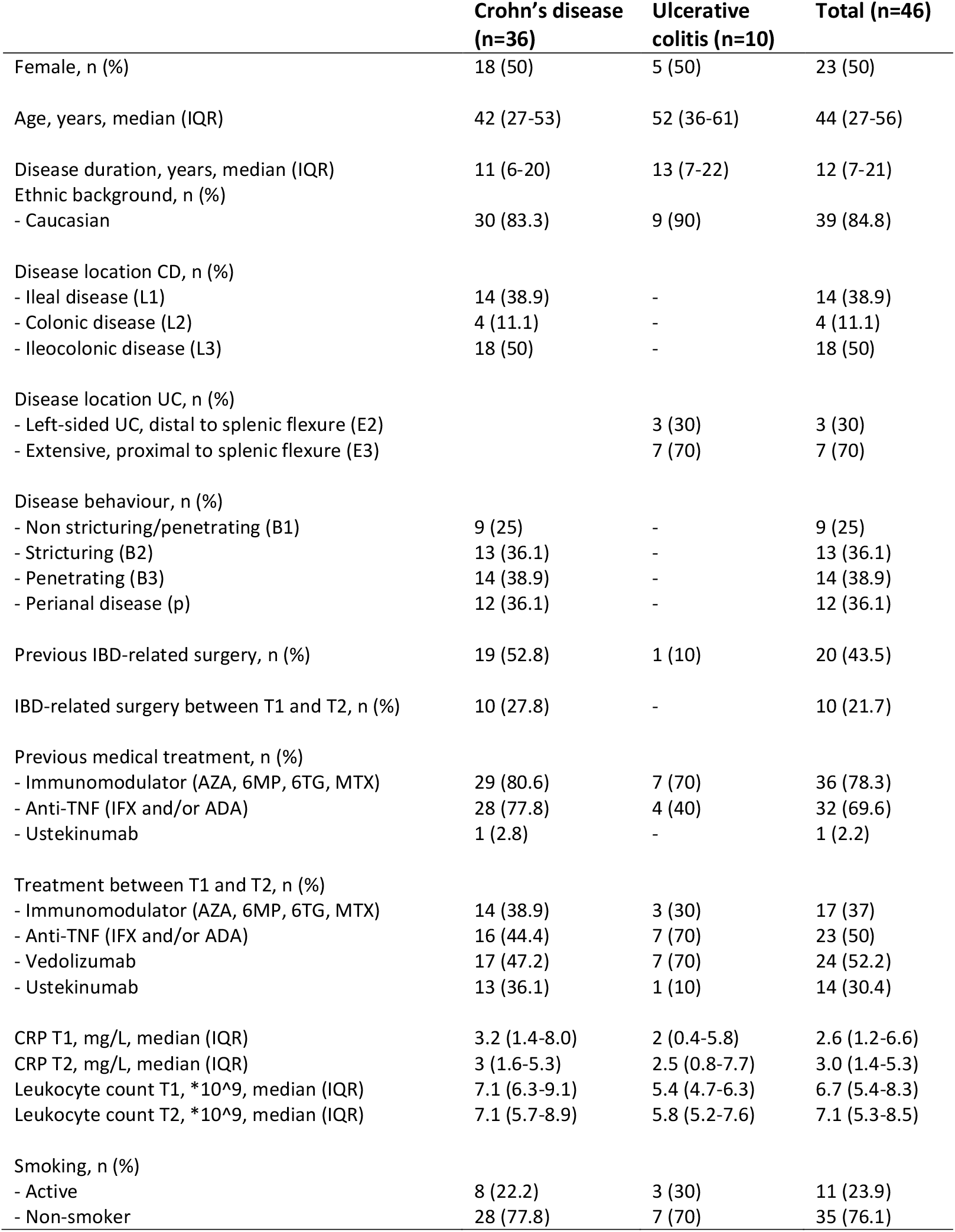
Baseline characteristics.

### Time-associated differential methylation expectedly associates with age-related CpGs

We first investigated the differences in methylation between both time points, identifying 194,391 (≈23%) differentially methylated positions (DMPs) when comparing T1 and T2 at a FDR-adjusted *p*-value of <0.05 (Figure 2a, b and c), which we termed time-associated DMPs. As our sample of interest was derived from peripheral blood, we investigated whether differences in the cellular composition were observable. Comparing the predicted blood cell composition yielded no significant differences (Figure 2d).

**Figure 2.**
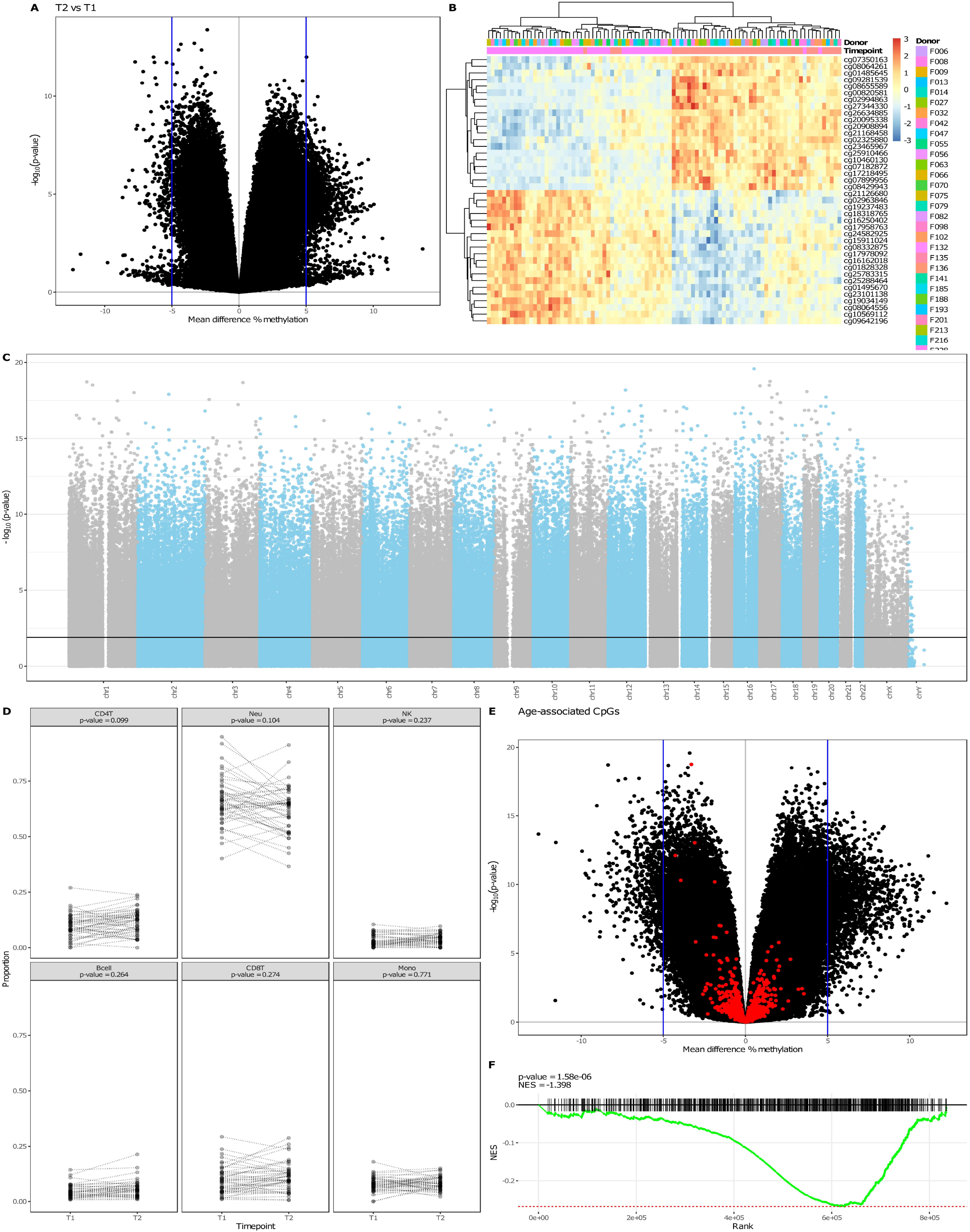
Time-variant methylated positions. A) A volcano plot depicting the mean difference in methylation between the two time points on the x-axis and the –log_10_(*p*-value) on the y-axis. B) Heatmap visualising the percentage methylation for the 25 most hyper- and 25 most hypo methylated DMPs. C) Manhattan plot showing the chromosomal distribution of all Illumina HumanMethylation EPIC array probes. Each dot represent a single CpG locus, dots above the black line are statistically significantly different between T1 and T2 (FDR-adjusted p-value ≤0.05). D) Estimated blood cell distribution stratified by time. Dashed lines connect samples obtained from the same donor. Statistical significance was calculated using a Mann-Whitney U test. E) Volcano plot colored for age-associated CpGs. F) Gene set enrichment analysis barcode plot representing the overrepresentation of the age-related CpGs among the time-associated DMPs.

By contrast, the time-associated differences were, expectedly, enriched for age-associated CpGs, which have been defined as the “epigenetic clocks” from Horvath^51^, Hannum^52^, Levine^53^, and Knight^54^ (Figure 2e). Furthermore, for these specific epigenetic clock CpGs, we observed a general hypomethylated pattern at T2 relative to T1 CpG sites (Figure 2f), suggesting that the observed differences in DNA methylation are enriched for age-related differences. Functional enrichment analyses of the time-associated DMPs displayed several cancer-associated pathways (Supplementary Figure 1).

### Time-invariant, stable methylated probes are enriched in genes involved in cell adhesion

In order to identify CpGs that were consistently methylated at both time-points, we performed intra-class correlation (ICC) analysis, which indicated that the majority of the CpGs (517.576 probes or around 60%) present poor intra-individual stability over time (ICC<0.50, Table 2). Conversely, 119.388 CpGs (≈14%) displayed a statistically significant high ICC (≥0.75), which we termed stably methylated positions (SMPs). We reasoned that probes that were associated with sites known to harbour genetic variants, both intentional and unintentional^59^, should present the highest stability as the genome of an individual typically does not change over the course of 7 years. Indeed, splitting the data by modality suggested that the CpG sites associated with known germline variants, namely those that were included for quality control purposes, presented high (>0.9) ICC values (Figure 3a and Table 2). Of the SMPs with ICC values over 0.9, 15.766 SMPs (around 2%) presented no indication that they bind predicted or potential genetic variants, which we classified as hyper-stable methylated positions (HSMPs, Table 3 and Figure 3b). Functional enrichment analyses of the SMPs and HSMPs indicated significant enrichment of genes involved in cell-cell signaling, adhesion and neurogenesis (Supplementary Figures 2 and 3).

**Table 2.**
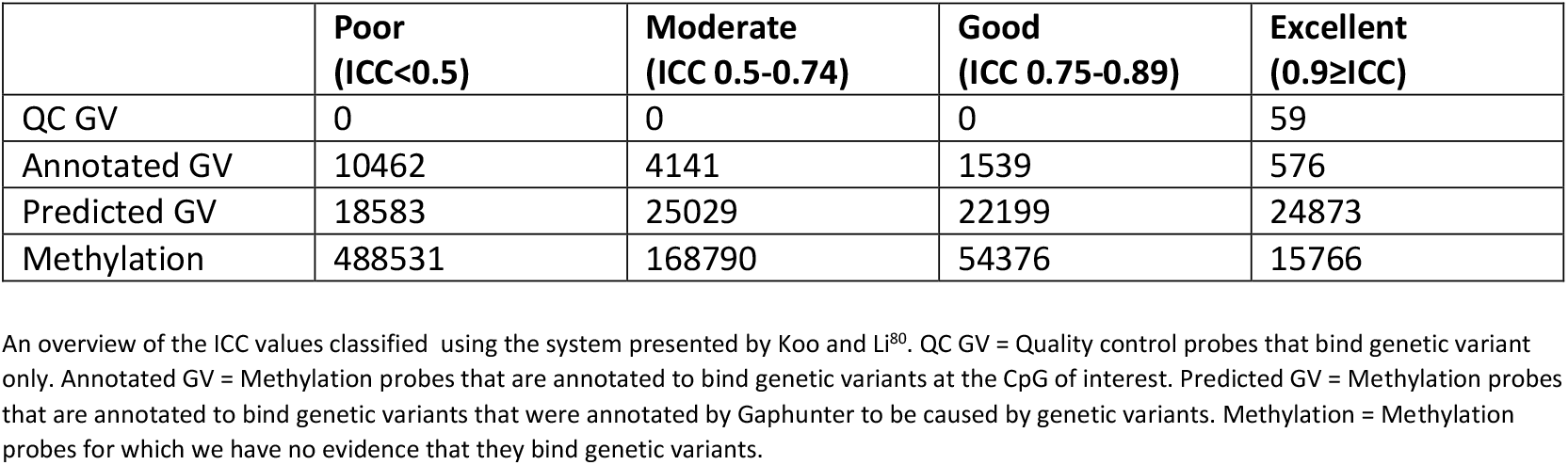
ICC scores.

**Figure 3.**
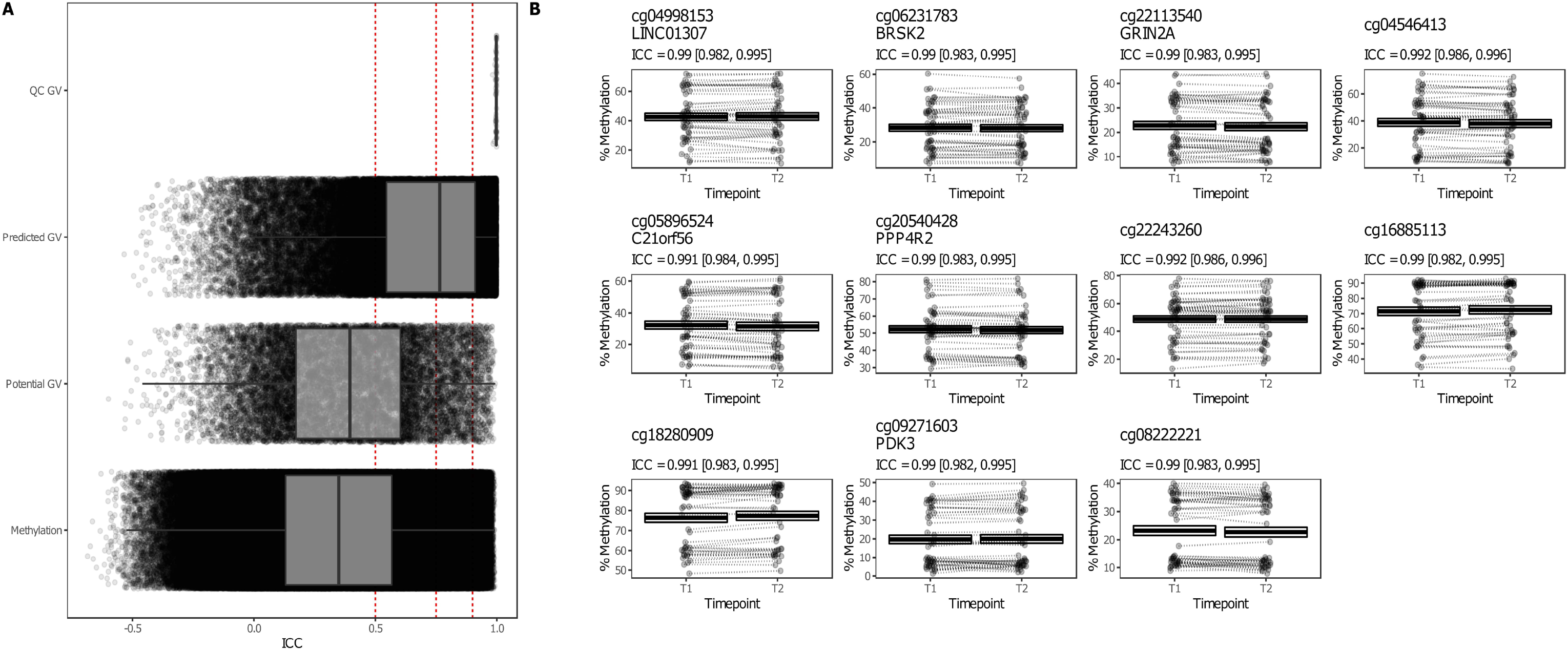
Time-invariant methylated positions. A) The intraclass correlation coefficients (ICC) stratified by probe type, with QC GV representing the aforementioned quality control probes that bind genetic variants (GV), the potential GV representing probes that were annotated with a genetic variant, and predicted GV representing probes that presented a methylation signal typically found when driven by a genetic variant. Red dashed lines represent the classification boundaries introduced by Koo and Li^80^, with blocks representing poor (ICC < 0.5), moderate (0.5 ≤ ICC < 0.75), good (0.75 ≤ ICC < 0.9), and excellent (0.9 ≥ ICC). B) Jittered visualisation of the 11 probes that present a time-invariant difference that is as stable as the aforementioned quality control probes with the percentage methylation on the y-axis and the time point on the x-axis. Dashed lines connect samples obtained from the same donor. The cross-bar visualisation represents the mean and standard error of the mean.

**Table 3.**
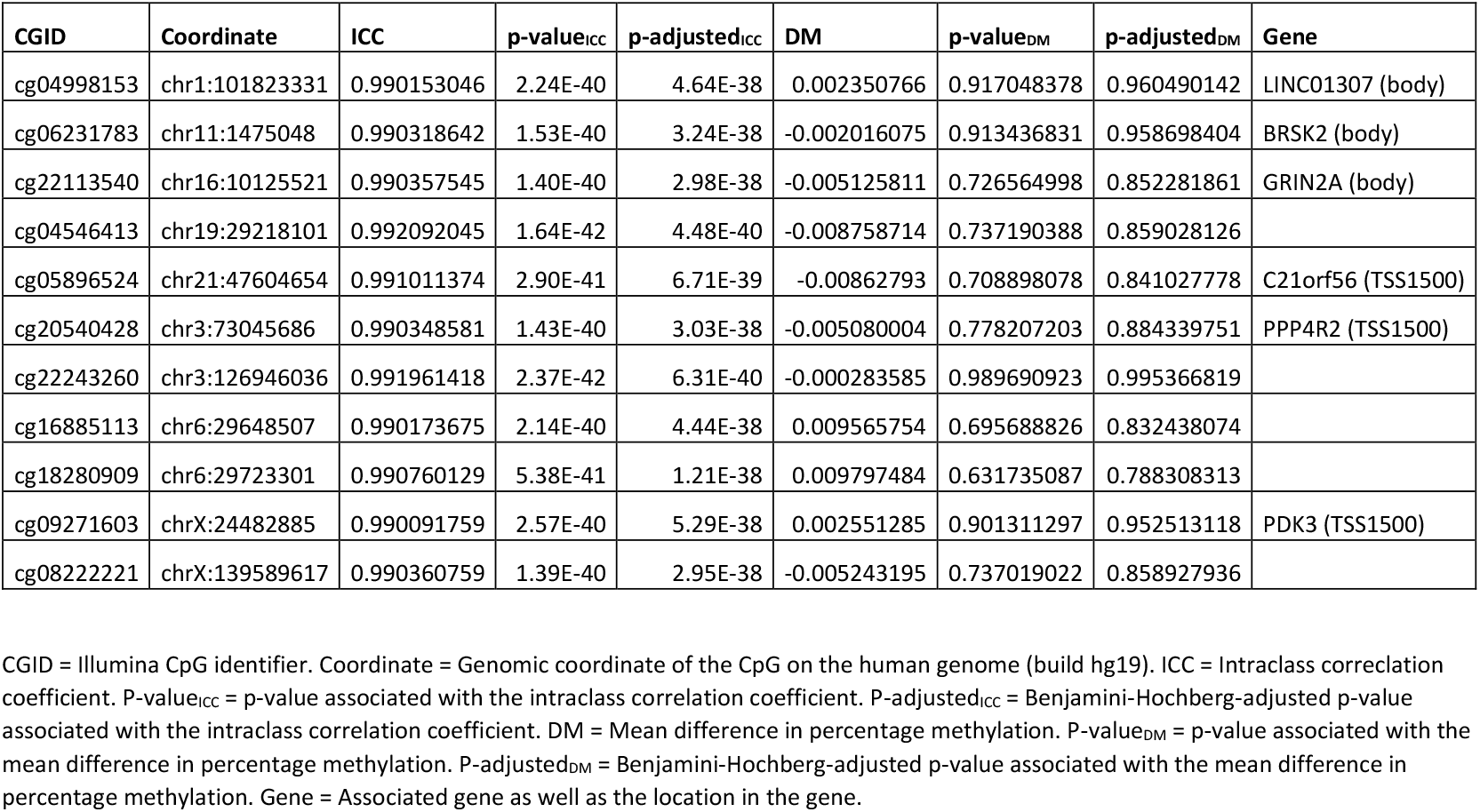
Hyper-stable methylated positions (ICC≥0.9)

### Stability analysis of previous IBD-associated DMPs, HLA and IBD-susceptibility genes

We next investigated whether previously reported IBD-associated DMPs were found to be invariant over time. To do so, we evaluated ICC values of 255 CD-associated, 103 UC-associated and 221 IBD-associated consistent DMPs identified in our systematic review and meta-analysis on CD-, UC-, and IBD-associated differential methylation^60^, which included a total of 552 samples (177 CD, 132UC and 243HC) from 4 different EWAS^14, 17–19^. Focusing on the stability of these DMPs in this cohort, we show that the majority (151 or 59.2% CD-associated, 73 or 70.9% UC-associated and 156 or 70.6% IBD-associated) present poor to moderate stability, indicating that the methylation status of these DMPs are affected by age or other exposures over time (Figure 4a, supplementary table 1). Nonetheless, 22 CD-associated (12.4%), 11 UC-associated (8.3%) and 24 IBD-associated loci (9.9%) show good to excellent stability over time, providing evidence that these CD-, UC-, or IBD-associated DMPs are unaffected by ageing or the exposures over time (Figure 4a, supplementary table 1).

**Figure 4.**
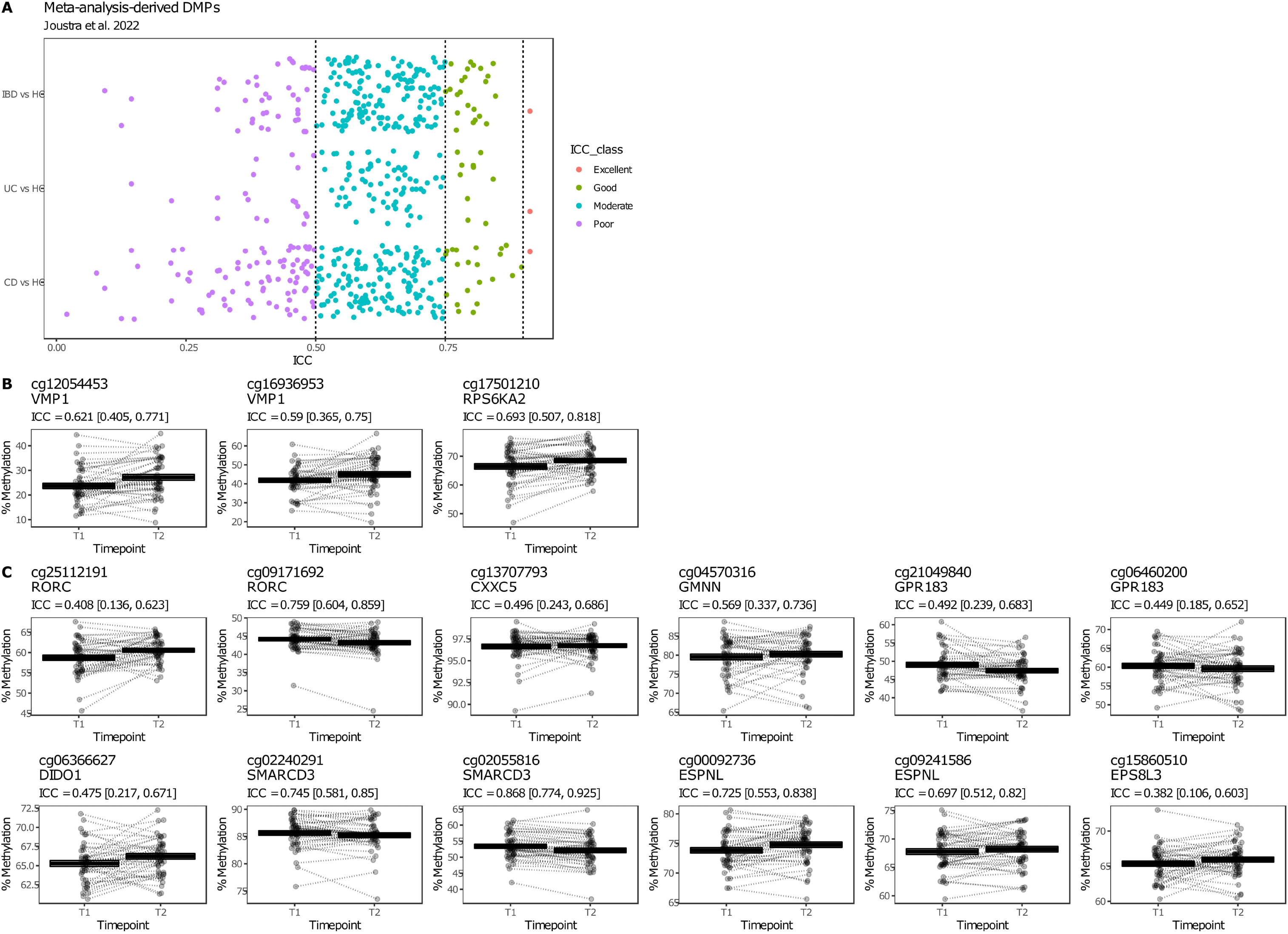
IBD-associated SMPs. A) The intraclass correlation coefficients (ICC) of consistent DMPs identified through meta-analyses of 4 EWAS^14^, ^17–19^ by Joustra et al.^60^ stratified by comparison (CD vs. HC, UC vs. HC and IBD vs. HC). Black dashed lines represent the classification boundaries introduced by Koo and Li^80^, with blocks representing poor (ICC < 0.5), moderate (0.5 ≤ ICC < 0.75), good (0.75 ≤ ICC < 0.9), and excellent (0.9 ≥ ICC). B) Jittered visualization of the IBD-associated DMPs as reported on by Adams et al.^18^, Ventham et al.^17^, as well as the CRP-independent probes reported on by C) Somineni et al.^23^ The percentage methylation is plotted on the y-axis and the time point on the x-axis. Dashed lines connect samples obtained from the same donor. The cross-bar visualization represents the mean and standard error of the mean.

Among the many IBD-associated DMPs, we specifically zoomed in on *VMP1* (cg12054453 and cg16936953) as well as *RPS6KA2* (cg17501210), as they were identified in multiple IBD-EWAS^17, 18, 23^ as well as shown found to be among the most significant IBD-associated DMPs in our meta-analysis^60^. We observed moderate consistency over time with noticeable overall hypermethylation at T2 relative to T1 for the aforementioned 3 CpGs (Figure 4b).

In addition to our meta-analysis, we also interrogated the CpGs that were CD-associated but CRP-independent as reported on by Somineni *et al*^23^. Interrogation thereof using our cohort revealed that cg25112191 (*RORC*), cg13707793 (*CXXC5*), cg21049840 (*GPR183*), cg06460200 (*GPR183*), cg06366627 (*DIDO1*), and cg15860510 (*ESP8L3)* presented poor consistency (ICC < 0.5), cg04570316 (*GMNN*),cg02240291 (*SMARCD3*), cg00092736 (*ESPNL*), and cg00092736 (*ESPNL)* presented moderate consistency (ICC 0.5-0.74), and cg09171692 (*RORC*) and cg02055816 (*SMARCD3*) presented good consistency (ICC 0.75-0.89) between both time points (Figure 4c).

We next interrogated the stability of all CpG loci associated with several well-known GWAS identified IBD risk genes involved in IBD pathogenesis, namely *ATG16L1, NOD2, IL23R, CARD9, FUT2, TYK2*, and *TNFSF15*^61, 62^, as well as specific IBD-associated major histocompatibility complex (MHC) encoding *HLA* genes previously reported on in GWAS studies, namely *HLA-DRB1, HLA-DQB1, HLA-DQA1, HLA-DPA1, HLA-DPB1, HLA-A, HLA-B*, and *HLA-C*^61, 63–66^. Comparing all IBD risk genes, we noticed that the *HLA* genes presented the highest stability, all of which had a median ICC score over 0.5 whereas the majority of CpGs that annotate to non-*HLA* IBD risk genes had poor ICC values (<0.5) (Figure. 5a). Nonetheless, for each of these non-HLA IBD risk genes, we identified highly stable methylated positions, several of which located to transcription start sites or 1^st^ exons (Supplementary figure 4 and supplementary table 2), implicating potential regulatory function.

**Figure 5.**
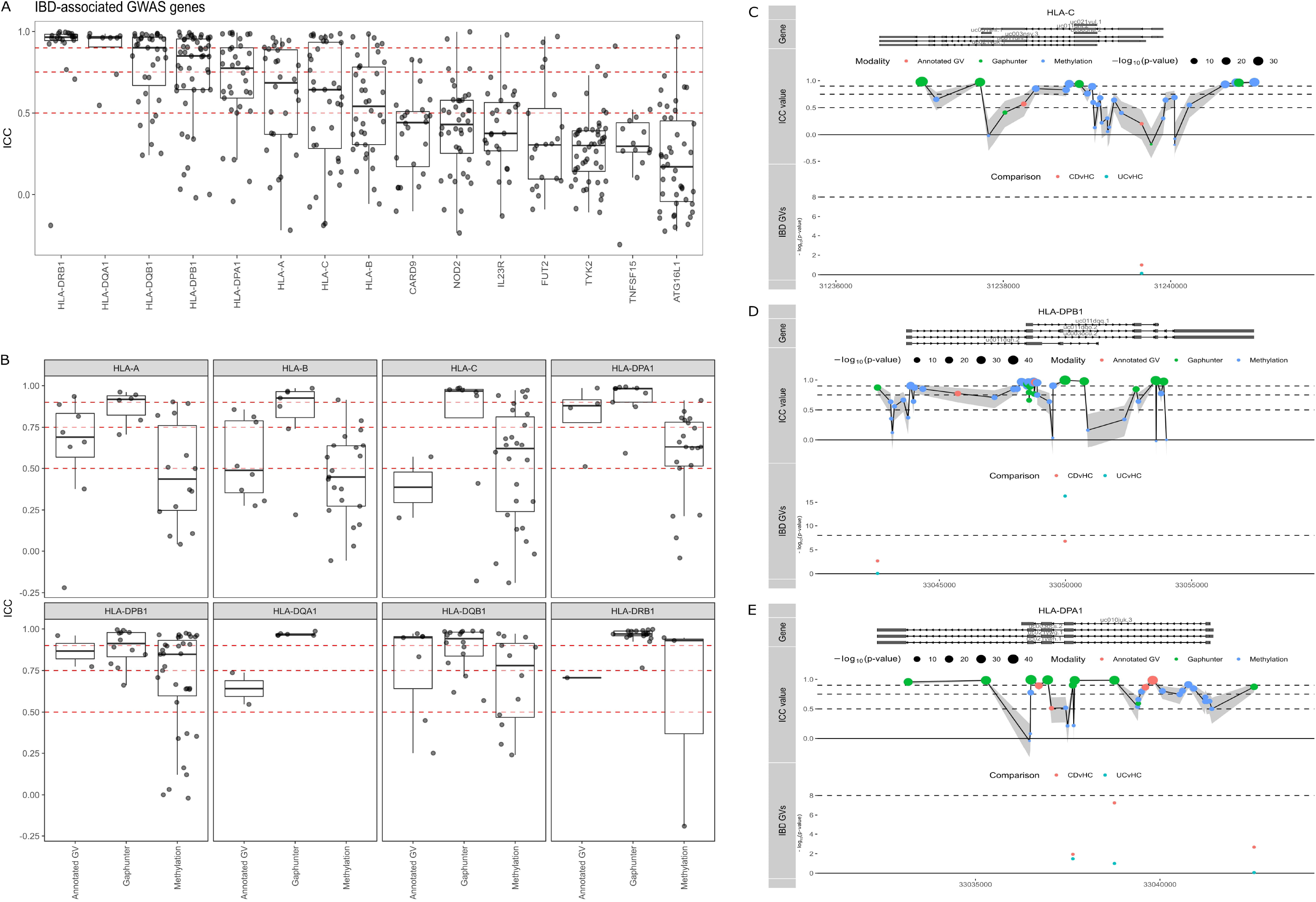
IBD risk genes. A) Visualisation of the intraclass correlation coefficients (ICC) for all CpGs annotated to IBD-associated GWAS genes. Boxplots show the overall median stability within each gene. Red dashed lines represent the classification boundaries introduced by Koo and Li^80^, with blocks representing poor (ICC < 0.5), moderate (0.5 ≤ ICC < 0.75), good (0.75 ≤ ICC < 0.9), and excellent (0.9 ≥ ICC). B) ICC values of all CpG loci of class I and II HLA genes. The potential GV representing probes that were annotated with a genetic variant, predicted GV representing probes that presented a methylation signal typically found when driven by a genetic variant, and methylation representing probes for which we have no evidence that they hybridise with any geneticm variant. Visualisations of the ICC values of all Illumina CpGs annotated to HLA-C (C), HLA-DPB1 (D) and HLA-DPA1 (E), relative to their position on each gene and grouped as potential GV (coral), predicted GV (green), or methylation (blue). Dots below represent known genetic variants as reported by Goyette^64^ for CD versus healthy controls (coral) and UC versus healthy controls (turquoise).

As DNA methylation measured using deamination technologies cannot distinguish DNA methylation from genetic variants located at the CpG of interest^59^, we investigated whether such technical artefacts were found among the *HLA* SMPs by interrogating the dbSNP (v151) database for catalogued variants, as well as by investigating for a typical clustered methylation signal when probes hybridise with genetic variants using Gaphunter^67^. Notably, most of the high ICC values were found for CpGs that presented some type of clustering typical of genetic variants, but were not necessarily catalogued in dbSNP (v151) (Figure. 5b). In addition, *HLA* class II genes (*HLA-DPA1, HLA-DPB1, HLA-DQA1, HLA-DQB1*, and *HLA-DRB1*) appeared to have a larger proportion of highly stable probes compared to *HLA* class I genes (*HLA-A*, *HLA-B*, and *HLA-C*) (Figure 5b). Besides technical artefacts, DNA methylation itself can be affected by genetic variants that occur in the vicinity^68^. As such, we cross-referenced our observations with a previous large-scale genotyping study of the *HLA* region in both CD and UC patients^64^. We indeed found multiple probes within the vicinity (<1Kb) of CD- or UC-associated *HLA* alleles, many of which did not appear to have an annotated dbSNP identifier, indicating that the observed differences were either unannotated GVs, or CpGs whose methylation status is strictly controlled by neighbouring IBD-associated GVs.

Notably, several highly stable probes (ICC ≥0.75) found to be annotated to *HLA-C, HLA-DPB1*, and *HLA-DPA1*, were located far away (>1Kb) from any of the IBD-associated GVs, did not associate with catalogued GVs, nor were identified as a potential GV by Gaphunter (Figure 5c, d and e, supplementary figure 5 and supplementary table 3), suggesting by default strong methylation stability over time.

## Discussion

Biomarker research often involves samples taken prior to or within a strictly pre-defined timeframe to the outcome of interest to mitigate the number of additional variables. In this study, we performed long-term longitudinal stability analyses of the PBL DNA methylome obtained from a cohort of adult IBD (36 CD and 10 UC) patients that were collected at two time points separated by a median of 7 years reflective of a real-life tertiary referral population.

Our observations indicate that the majority of all loci (≈60%) measured on the Illumina HumanMethylation EPIC BeadChip array present notable intra-individual variation in methylation over time (ICC <0.5), which is enriched for age-associated CpGs. Nonetheless, not all time-associated DMPs in our cohort were previously reported as age-associated CpGs. While we observed no significant differences in CRP, leukocyte count, and estimated cellular composition between both time points, other external or environmental exposures, such as smoking, dietary alterations, therapy failure or switch, IBD-related surgery or disease progression might have altered methylation status contributing to the observed time-associated differences. By contrast, 119.388 (≈14%) and 41.274 (≈5%) loci presented a highly stable pattern across both time points with ICC values ≥0.75 and ≥0.90 respectively. Such loci retained their degree of methylation even after the aforementioned known IBD-associated and unknown external exposures, suggesting time-stability.

Previous studies investigating DNA methylation stability in PBL of healthy adults using both the Illumina HumanMethylation EPIC BeadChip array, as well as its predecessor the Illumina HumanMethylation 450k BeadChip array, presented similar observations. In these studies, 16.9 – 23% of the CpG loci presented a moderate/good (ICC 0.50 - 0.79) while 8.3-12.9% of the CpG loci presented a good/excellent (ICC ≥0.8) stability over a span of 1 to 6 years^39, 41^. Focusing on IBD, when interrogating 255 CD-associated, 103 UC-associated and 221 IBD-associated DMPs identified in our own meta-analysis^60^ of 4 IBD EWAS^14, 17–19^, we observed that the majority presented poor to moderate stability, suggesting that the aforementioned IBD-associated loci might also be affected by exposures over time that might or might not be related with IBD. While interesting, such time-variant probes should be interpreted with care when used as predictive biomarkers given their association with exposures that occurred during both time points. To that end, our data could be used as a resource to preselect time-invariant CpG loci before independent validation when performing IBD-associated EWAS^69^ thereby increasing the potential to identify replicable predictive biomarkers better reflecting the underlying biology of IBD. In addition, such an approach would enable a larger pool of samples to be used as samples need not to be obtained within the same age range when performing DNA methylation studies on IBD and its phenotypes.

When specifically interrogating the IBD-associated probes cg12054453 (*VMP1*), cg16936953 (*VMP1*), and cg17501210 (*RPS6KA2*), we find moderate consistency over time with a noticeable hypermethylation at T2 compared to T1. Similar analyses performed on the CD-associated yet CRP-independent probes reported by Somineni et al.^23^ showed that the majority of these CpGs did not present long-term stability in our cohort. Differences between our observations and that of Somineni might be attributable to differences between adult and paediatric cohorts, or might simply reflect non-inflammatory changes in methylation that occur over time. By contrast, cg09171692 (*RORC*) and cg02055816 (*SMARCD3*) presented high ICC values (0.76 and 0.87 respectively) indicating good stability in our cohort, irrespective of CRP or non-inflammatory exposures. Notably, both genes have previously been associated with IBD in multiple studies^23, 70, 71^ and therefore show promise as stable IBD-associated loci.

Given the complex, multifactorial nature of IBD^2^, focussing on the interplay between genetic variation and DNA methylation rather than single gene mutations alone might prove more useful in understanding its molecular aetiology. Previous EWAS of mucosal tissue^72^ and peripheral blood^73^ both demonstrated differential methylation between IBD and controls for well-known GWAS identified IBD-associated risk genes, suggesting differential methylation of key risk genes to affect disease susceptibility. Our observations corroborate this hypothesis, showing highly stable methylation for particular CpG loci within these risk genes. Interestingly, several of these HSMPs were located in or near to the transcription start sites, potentially regulating gene transcription by maintaining the aberrant phenotype^18^.

There has been extensive interest in (epi)genetic alterations of the highly polymorphic *HLA* region related to IBD pathogenesis, most consistently reported for *HLA* class II genes involved in the presentation of bacterial antigens to CD4+ T-cells^14, 18, 61, 63–66, 74, 75^. Specifically, genetic variation of classical *HLA* genes has been suggested to play a role in the aberrant response to the dysbiotic microbiome observed in IBD^63^ with particular impacts for the response to biological treatment^76–78^ and the formation of anti-drug antibodies^66^. However, translation of the results into clinical practice has proven to be difficult due to the high number of polymorphisms of HLA αβ heterodimers and strong linkage disequilibrium^63^.

In our study, we observed multiple HSMPs in HLA genes, suggesting that the DNA methylation profile of these genes is very stable over time. Our results corroborate with previous array data showing highly significant correlations between CpG loci on several *HLA* class II genes of neonates compared to toddlers (r=0.83) and adults (r=0.88) with type 1 diabetes^79^. While further interrogation of these HSMPs indicates that multiple CpGs might be actual genetic variants, we also find multiple HSMPs that are not genetic variants. Nonetheless, several of such epigenetic HSMPs do occur within the vicinity of known IBD-associated HLA-variants, providing evidence that particular HLA-alleles might impart a strong, stabilising effect on the epigenome.

### Strengths and limitations

To our knowledge, we are the first to assess the stability of the DNA methylome obtained from PBL of IBD patients with a median 7 year follow-up period in a real-life disease exposure setting. This study is explorative in nature, using a moderate sample size without prior power calculation. Nonetheless, we note that studies of similar design have been conducted with a similar sample size^39, 42, 69^. While we can be reasonably confident in identifying the time-invariant aspect of the SMPs, we cannot fully eliminate the possibility that the SMPs would remain stable in a more diverse IBD cohort, as the typical markers of inflammation (CRP and leukocyte count) and cellular distribution were hardly different between both time points.

## Conclusion

We observe considerable variability in DNA methylation measurements taken from PBL at two different time points separated by a median of 7 years. By contrast, around 14% of all CpG loci could be considered highly stable even after IBD-specific exposures during the two points. Focussing on these CpG loci during biomarker discovery might result in the identification of biologically relevant and more robust IBD-associated epigenetic biomarkers with an increased probability of independent replication.

---

**Supplementary Figure 1.** Functional enrichments analyses using GO-term and KEGG pathways for the time-associated DMPs

**Supplementary Figure 2.** Functional enrichments analyses using GO-term and KEGG pathways for the stable methylated positions (SMPs)

**Supplementary Figure 3.** Functional enrichments analyses using GO-term and KEGG pathways for the hyper-stable methylated positions (HSMPs).

**Supplementary Figure 4.** Visualisation of the ICC values of all Illumina CpGs annotated to IBD GWAS risk genes, relative to their position on each gene. Dots represent individual CpG loci, dots above the red dashed line are considered SMPs (ICC≥0.75) whereas dots above the blue dashed line are HSMPs (ICC≥0.9).

**Supplementary Figure 5.** Visualisation of the ICC values of all Illumina CpGs annotated to HLA-A (A), HLA-B (B), HLA-DQA1 (C), HLA-DQB1 (D) and HLA-DRB1 (E) relative to their position on each gene and grouped as potential GV (coral), predicted GV (green) or methylation (blue). Dots below represent known genetic variants as reported by Goyette^64^ for CD versus healthy controls (coral) and UC versus healthy controls (turquoise).

**Supplementary Table 1.** Summary results of the ICC results for each of the identified consistent DMPs by Joustra et al.^60^, stratified by comparison (CD vs. HC, UC vs. HC and IBD vs. HC).

**Supplementary Table 2.** Summary of the ICC results for each of the CpGs located on the IBD GWAS risk genes.

**Supplementary Table 3.** Summary of the ICC results for each of the CpGs located on the HLA genes.

