## Supplementary figures and images for "Long-term temporal stability of peripheral blood DNA methylation alterations in patients with inflammatory bowel disease"

### Supplementary figure 1

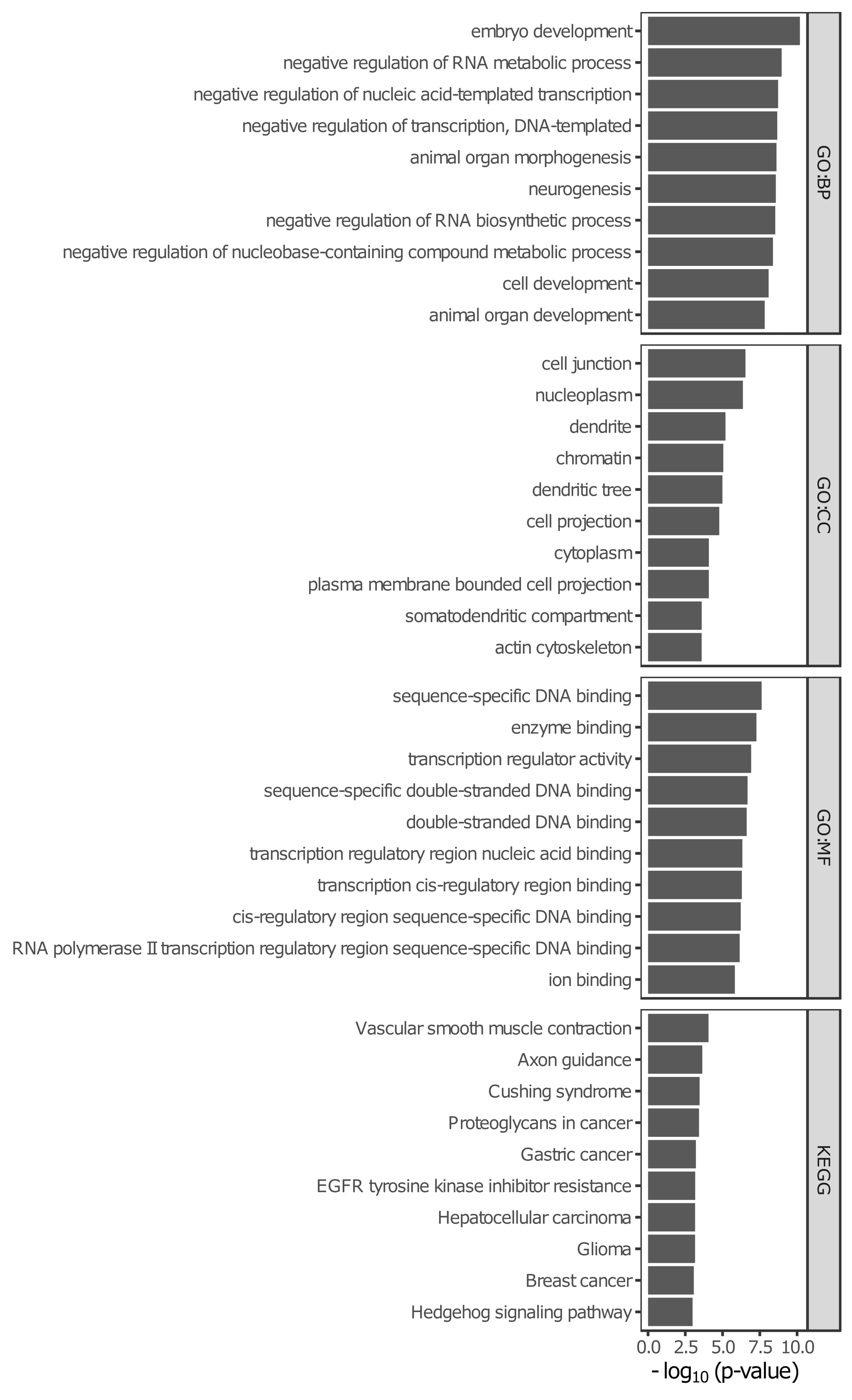

### Supplementary figure 2

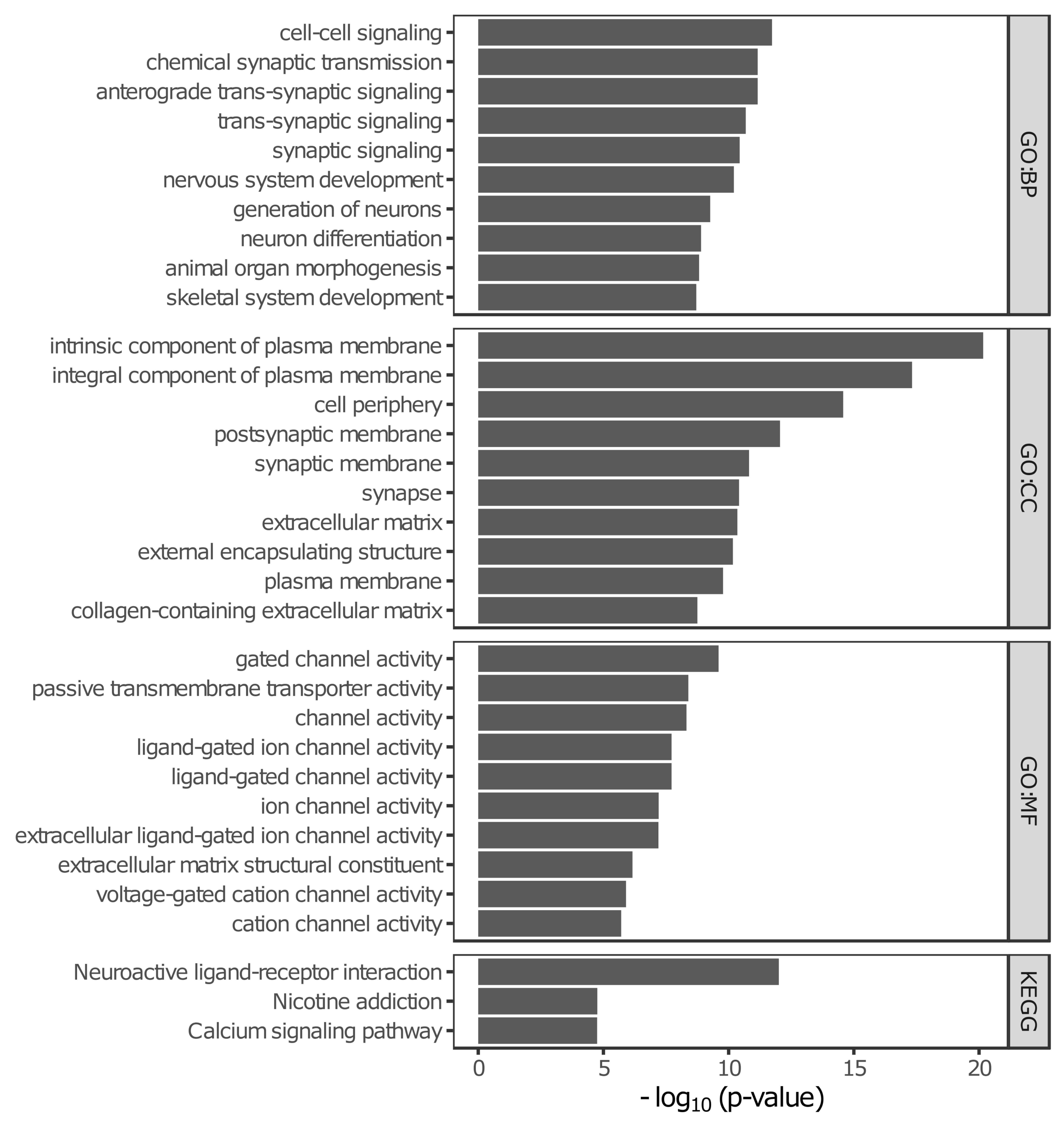

### Supplementary figure 3

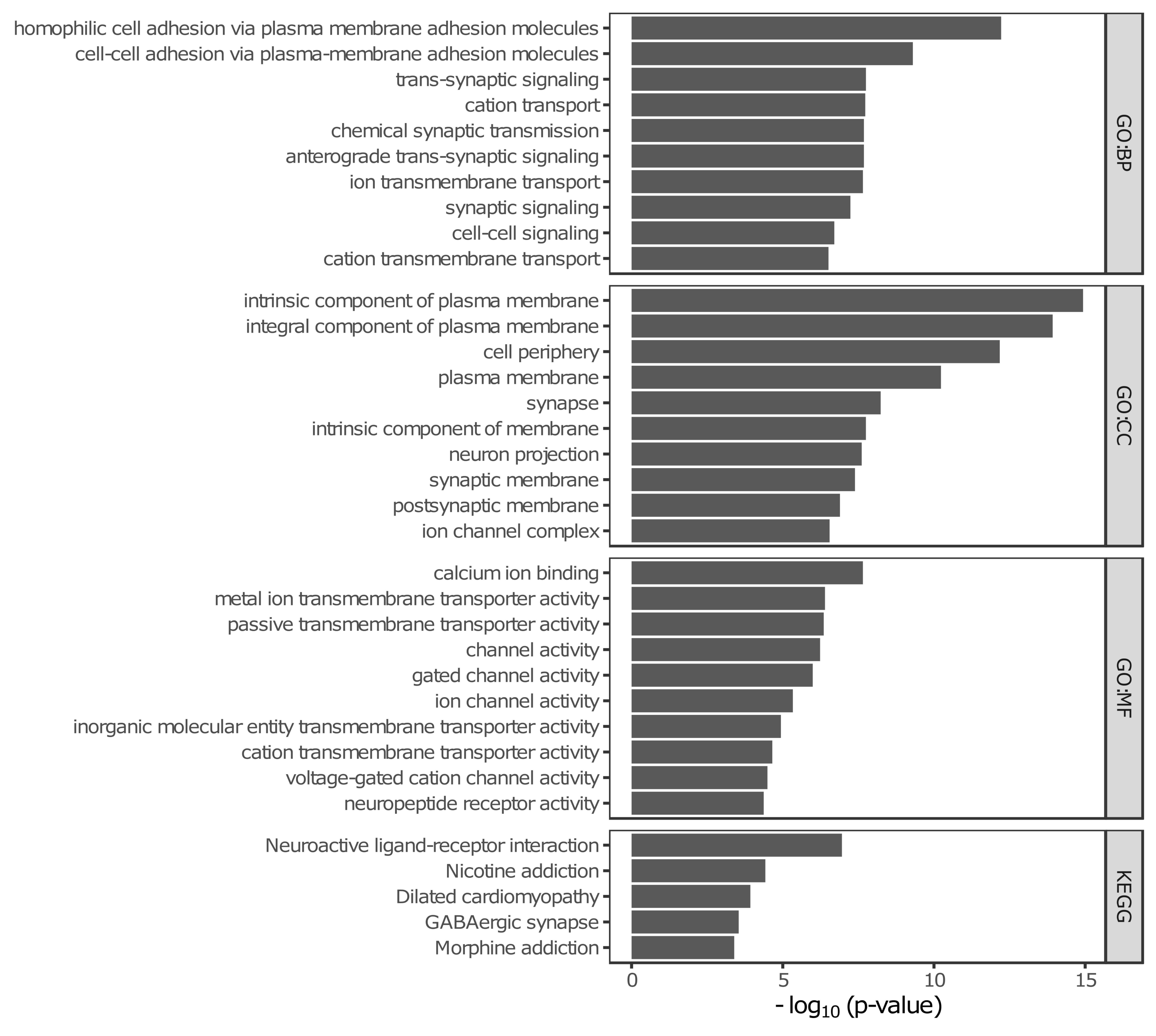

### Supplementary figure 4

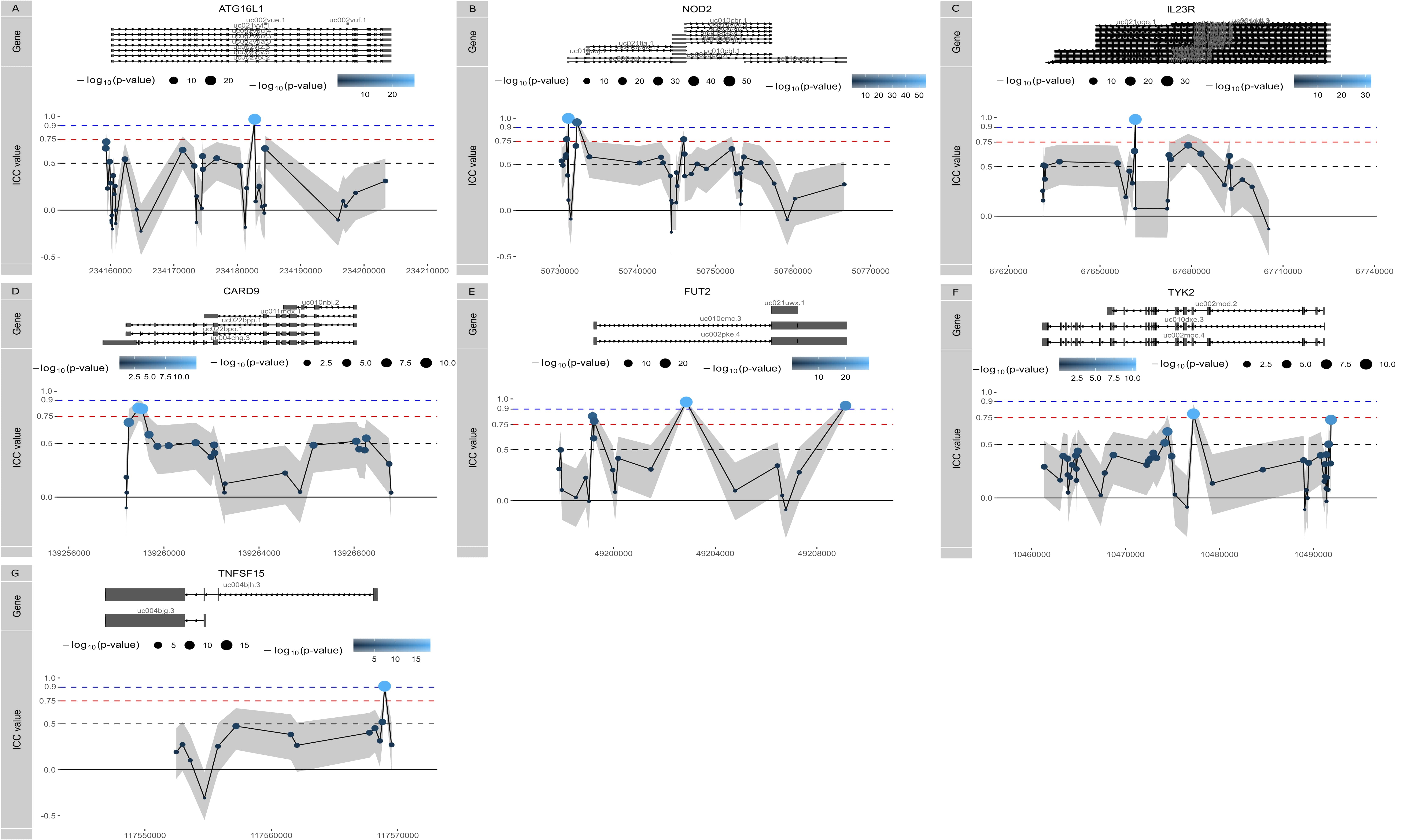

### Supplementary figure 5

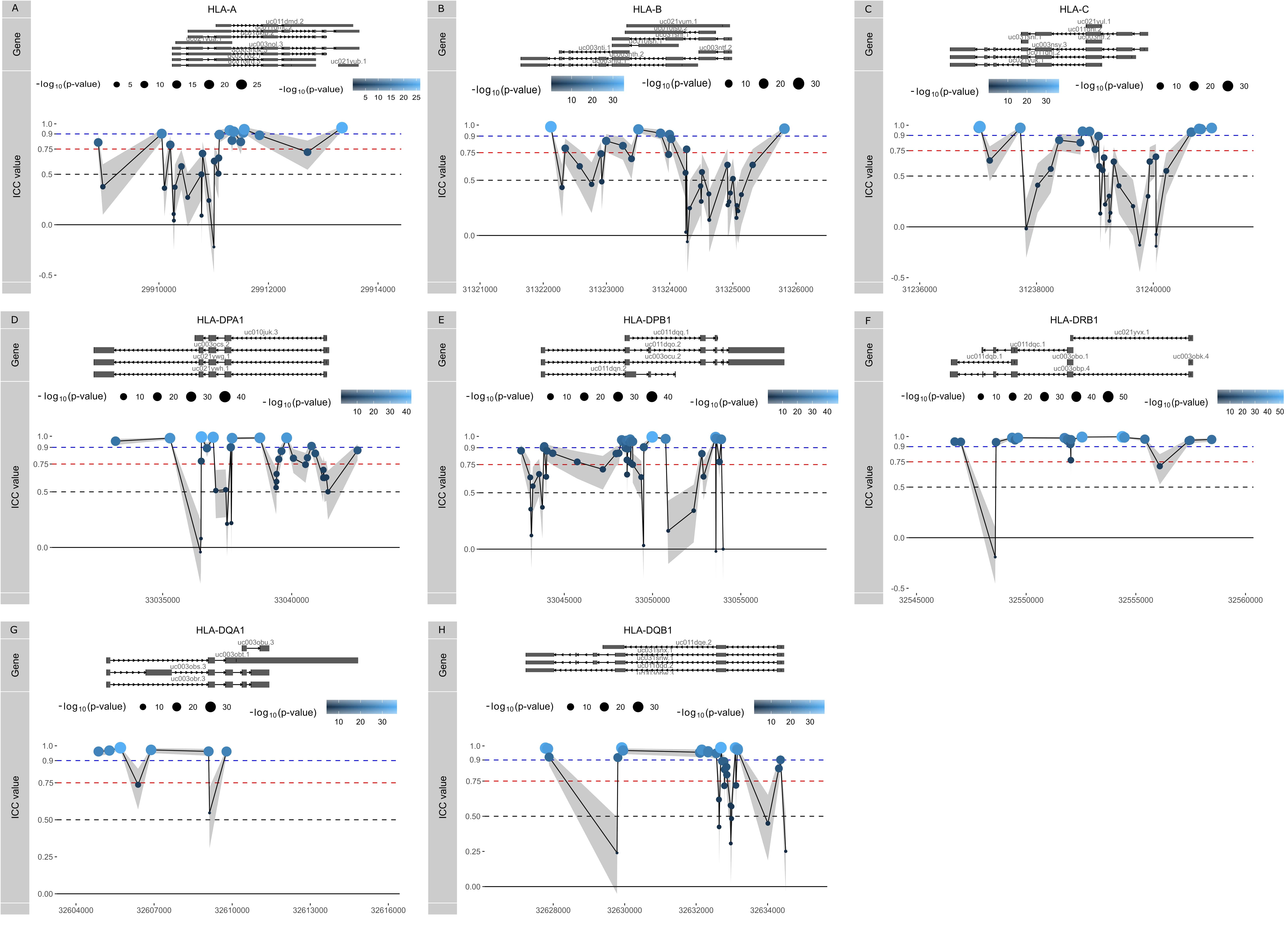
